# Water immunity overrides stomatal immunity in plant resistance to *Pseudomonas syringae*

**DOI:** 10.1101/2024.10.07.617076

**Authors:** Jasmin Kemppinen, Maximillian Pollmeier, Sanna Ehonen, Mikael Brosché, Maija Sierla

## Abstract

Stomata play crucial roles in the multilayered defense system against pathogens. Upon pathogen perception, stomata close promptly, establishing the first line of defense known as stomatal immunity. The bacterial pathogen *Pseudomonas syringae* (*Pst*) exploits open stomata for entry, however, it can also induce stomatal closure at post-invasive stages to enhance apoplastic hydration. This creates a favorable environment for *Pst* proliferation, evident as water-soaked lesions on leaves. During the post-invasive stages of *Pst* infection, plants deploy a second layer of stomatal defense by reopening their stomata, a process termed water immunity. To evaluate the relative importance of stomatal versus water immunity, we utilized a diverse set of Arabidopsis (*Arabidopsis thaliana*) mutants with impaired stomatal function and monitored bacterial growth, stomatal behavior, and water-soaking capacity after *Pst* pv. *tomato* DC3000 infection. Most mutants with constitutively open stomata and disrupted stomatal closure were more resistant against *Pst* than wild-type plants. Also, while some stomatal mutants displayed similar stomatal behavior at the initial steps of defense, their disease outcomes were the opposite, suggesting that stomatal immunity is not the determining factor in disease resistance. Instead, we discovered that the water-soaking capacity, associated with stomatal status at later stages of infection, i.e., water immunity, dictates the disease outcome. Our results show that water immunity can override the lack of stomatal immunity in plant resistance to *Pst*. We also address previous discrepancies in literature showing contradicting results for pathogen growth on stomatal mutants, highlighting the challenges in dissecting stomatal effects on plant resistance.

## Introduction

Stomata are pores that cover aerial plant parts and function as integral sites for gas exchange. Flanking guard cells open or close the pores in response to environmental cues, balancing CO_2_ intake and water loss. Open stomata also serve as entry points for harmful organisms. Plants can restrict pathogen entry by rapidly closing stomata upon detecting pathogen-associated molecular patterns (PAMPs), a response known as stomatal immunity (Melotto et al., 2006). PAMP perception by receptors trigger signaling cascades involving intracellular kinases, calcium fluxes and reactive oxygen species (ROS) burst to activate anion channels, leading to depolarization of guard cell membranes, and stomatal closure (Melotto et al., 2024). One of the most studied model pathogens that use stomata as entry point is *Pseudomonas syringae* pv. tomato DC3000 (*Pst*). Guard cells recognize *Pst* by its flagellin epitope flg22, which is bound by the cell-membrane receptor complex formed by FLAGELLIN-SENSITIVE2 (FLS2), BRI1-ASSOCIATED KINASE1 (BAK1), and BOTRYTIS-INDUCED KINASE1 (BIK1).

Upon activation, the signaling module initiates stomatal closure in stomatal immunity (Figure 1A; Gómez-Gómez & Boller, 2000; Chinchilla et al., 2007; Lu et al., 2010). *Pst* can further alter guard cell responses by secreting coronatine (COR), a jasmonic acid mimic, to reopen stomata after the initial recognition to promote entry into the leaf tissues, thus bypassing stomatal defenses (Melotto et al., 2006).

**Figure 1.**
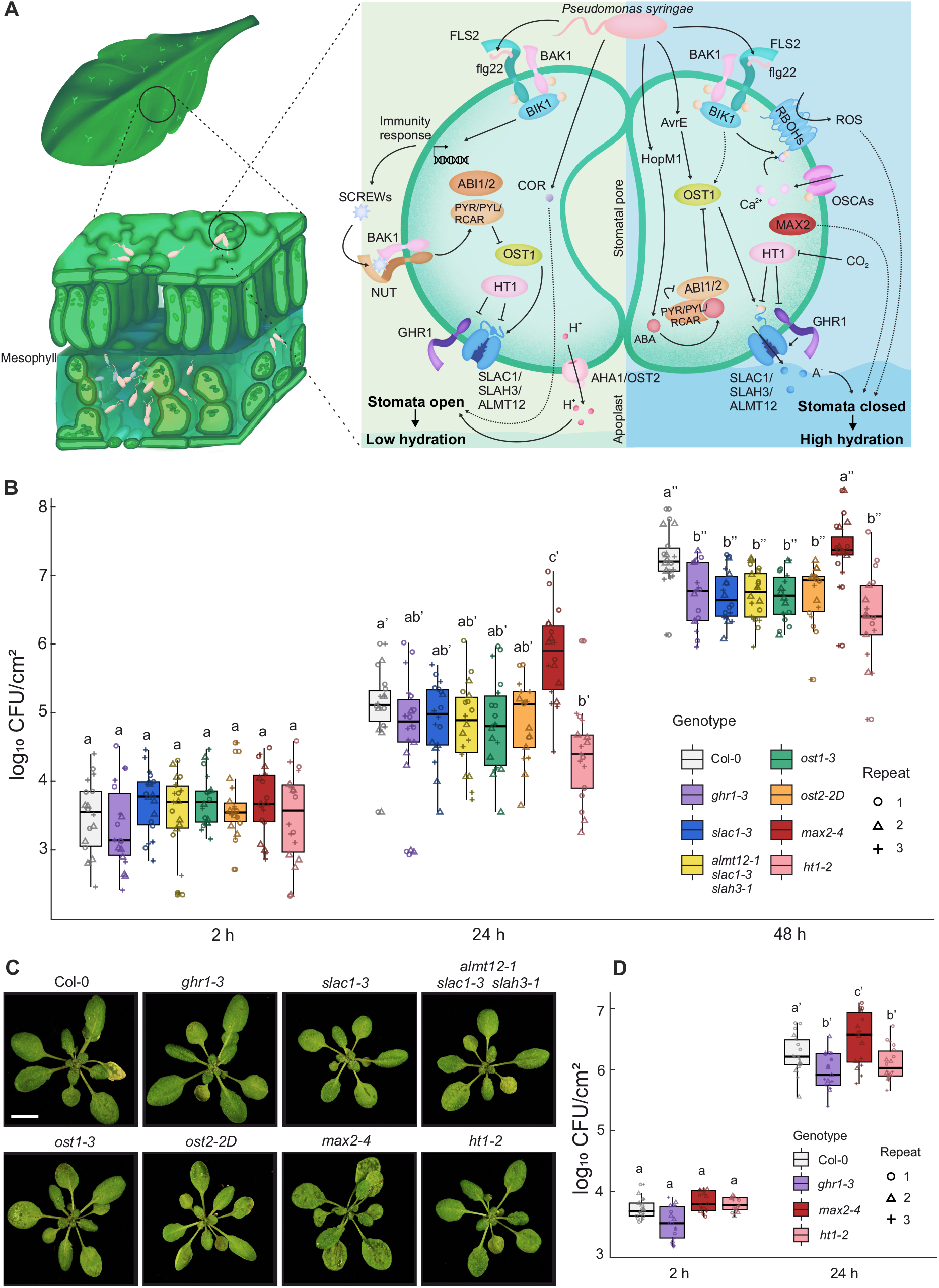
Signaling pathways that regulate stomatal movements during *Pseudomonas syringae* (*Pst*) infection and *Pst* growth in Arabidopsis mutants with stomatal dysfunction phenotypes. **A)** Apoplastic hydration in the mesophyll is crucial for *Pseudomonas syringae* (*Pst*) proliferation and requires closed stomata. *Pst* enhances its virulence by manipulating guard cell signaling, initially promoting stomatal opening to facilitate entry during early infection (via COR and JA signaling) and later driving stomatal closure to induce water-soaking (via enhanced ABA signaling). Plants defend themselves by regulating stomatal aperture at appropriate times during infection; initial stomatal closure restricts *Pst* entry and reopening at post-invasive stages prevents apoplastic hydration. Stomatal opening involves suppression of closure pathways, leading to membrane hyperpolarization through proton pumping (AHA1/OST2) and activation of inward-rectifying potassium channels, driving osmotic flow into guard cells. Conversely, stomatal closure requires activation of SLAC1, SLAH3 and ALMT12 anion channels, leading to membrane depolarization and activation of outward-rectifying potassium channels, followed by water efflux from guard cells and stomatal closure. Numerous other components are required to process environmental and hormonal signals, and include ABA perception, kinases such as OST1, HT1, BIK1, and secondary messengers, including ROS and calcium fluxes. Other guard cell signaling components have less defined function and include the pseudokinase GHR1 and components of E3 ubiquitin ligase complexes, such as MAX2. For detailed description of the main components and pathways, see the main text. Abbreviations: ABA, abscisic acid; ABI1/2, ABA-INSENSITIVE1/2; AHA1/OST2, AUTOINHIBITED H+ -ATPASE/OPEN STOMATA2; ALMT12, ALUMINUM-ACTIVATED MALATE TRANSPORTER12; BAK1, BRI1-ASSOCIATED KINASE1; BIK1, BOTRYTIS-INDUCED KINASE1; COR, coronatine; FLS2, FLAGELLIN-SENSITIVE2; GHR1, GUARD CELL HYDROGEN PEROXIDE-RESISTANT1; HT1, HIGH LEAF TEMPERATURE1; MAX2, MORE AXILLARY GROWTH2; NUT, PLANT SCREW UNRESPONSIVE RECEPTOR; OSCA, REDUCED HYPEROSMOLALITY-INDUCED Ca2+ INCREASE; OST1, OPEN STOMATA 1; PYR/PYL/RCAR, PYRABACTIN RESISTANCE1/ PYR1-LIKE/REGULATORY COMPONENTS OF ABA RECEPTORS; RBOH, RESPIRATORY BURST OXIDASE HOMOLOG; ROS, reactive oxygen species; SCREW, SMALL PHYTOCYTOKINES REGULATING DEFENSE AND WATER LOSS; SLAC1, SLOW ANION CHANNEL1; SLAH3, SLAC1 HOMOLOGUE3. **B)** Bacterial titers in Arabidopsis mutants with stomatal dysfunction phenotypes. Plants were spray-infected with *Pst* (OD_600_= 0.2 or 1 ×10^8^ CFU/mL) and bacterial titers were quantified 2 h, 24 h and 48 h post-infection. **C)** Visual symptoms of *Pst* spray-inoculated Arabidopsis rosettes 48 h post-infection. Scale bar = 10 mm. **D)** *Pst* growth in Col-0, *ghr1-3, max2-4* and *ht1-2* after syringe infiltration. Three leaves from each plant were infiltrated with *Pst* (OD_600_= 0.02 or 1 ×10^7^ CFU/mL), and bacterial titers were quantified 2 h and 24 h after infiltration. In B) and D), colony-forming units (CFU/cm^2^) were calculated from bacterial titers extracted from three leaves per plant. Box plots represent three independent repeats (n = 18 per genotype), shown with medians and interquartile ranges. Whiskers represent the range of data within 1.5 times the interquartile range from the first and third quartiles. Log-transformed CFU counts are displayed with different shapes indicating each repeat. A linear mixed-effects model followed by Tukey’s posthoc test was applied to assess statistical differences within each time point (indicated by an apostrophe). Genotypes sharing the same letter are not significantly different.

*Pst* can also induce stomatal closure at later stages of infection to form an aqueous phase inside the apoplastic spaces of the mesophyll (Xin et al., 2016). *Pst* use type III effectors, HopM1 and AvrE1, to promote abscisic acid (ABA) signaling and stomatal closure (Hu et al., 2022; Roussin-Léveillée et al., 2022). *Pst* AvrE-family effectors can form water-permeable pores on plant cell membranes, increasing the hydration in mesophyll by releasing cell contents (Nomura et al., 2023). These factors contribute to hydration of the apoplast, observed as water-soaking lesions. To counter this post-invasive virulence mechanism, plants employ a secondary, temporally regulated defense strategy known as water immunity, also referred to as apoplast or mesophyll immunity (Hou et al., 2024; Jian et al., 2024). In water immunity stomatal reopening is facilitated by plant-secreted peptides SMALL PHYTOCYTOKINES REGULATING DEFENSE AND WATER LOSS (SCREWs), as well as light-mediated salicylic acid (SA) signaling, both acting antagonistically to ABA signaling (Liu et al., 2022; Lajeunesse et al., 2023).

Early defense through stomatal immunity is considered crucial in plant defense against *Pst* and is thought to be an important determinant for disease outcomes. Studies on stomatal mutants show increased susceptibility to *Pst* due to disrupted stomatal immunity, leading to higher pathogen entry (Supplementary Table S1). However, recent research focusing on water immunity has shown that *Pst*-induced aqueous phase is prevented when stomata are open (Lajeunesse et al., 2023). Depending on the research angle, many articles, although studying the same Arabidopsis (*Arabidopsis* thaliana) genes required for stomatal regulation, show contradictory results regarding bacterial growth (Supplementary Table S1). For example, OPEN STOMATA1 (OST1) mutant *ost1-2* shows heightened susceptibility after vacuum-infiltration (Melotto et al., 2006), whereas *ost1-3* exhibits wildtype-like susceptibility after dip-inoculation (Jalakas et al., 2017; Ou et al., 2022). Similarly, OPEN STOMATA2 (OST2) mutant *ost2-2d* was found to be more susceptible against *Pst* after spray-infection, while exhibiting similar resistance as Col-0 after syringe-infiltration, emphasizing the role of stomatal immunity (Zhou et al., 2015).

In the light of recent insights into water immunity and its importance for later stages of *Pst* colonization, we set out to examine which layer of immunity, stomatal immunity, or water immunity, is more important for the success of *Pst* infection. We used mutants with constitutively open stomata or with disruptions in stomatal closure, including mutants of anion channels SLOW ANION CHANNEL1 (SLAC1), SLAC1 HOMOLOG3 (SLAH3), and ALUMINUM-ACTIVATED MALATE TRANSPORTER12 (ALMT12) (*slac1-3* and the *almt12-1 slah3-1 slac1-3* triple mutant; Vahisalu et al., 2008; Jalakas et al., 2021), ABA signaling component OST1 (*ost1-3/srk2e*; Yoshida et al., 2002), receptor-like pseudokinase GUARD CELL HYDROGEN PEROXIDE-RESISTANT1 (GHR1; *ghr1-3*; Sierla et al., 2018), F-box protein MORE AXILLARY GROWTH2 (MAX2; *max2-4*; Piisilä et al., 2015) and a mutant with overly active H^+^-ATPase OST2 and stomatal opening (*ost2-2D*; Merlot et al., 2007) (Figure 1A; Supplemental Table S2). We also included a mutant of HIGH LEAF TEMPERATURE1 (HT1) with constitutively closed stomata (*ht1-2;* Hashimoto et al., 2006). We observed that despite allowing greater pathogen entry through more open stomata, most of the stomatal mutants showed enhanced resistance suggesting that water immunity overrides impaired stomatal immunity.

## Results

### Most mutants with impaired stomatal function show resistance to *Pst*

As previous literature showed discrepancies in *Pst* growth on stomatal mutants (Supplemental Table S1), we selected a broad set of mutants with impaired stomatal responses for *Pst* growth assays to test plant resistance. We spray infected 3.5-week-old Arabidopsis plants with *Pst* and followed bacterial growth and disease symptom development for two days (Figure 1, B and C). All mutants exhibited less bacterial growth than Col-0 except for *max2-4*, which was highly susceptible against *Pst*.

The contribution of initial stomatal openness to bacterial infection can be estimated by comparing spray infection (where bacteria enter through stomata) with syringe infiltration (where bacteria are directly injected to the interior of the leaf). Syringe infiltration thus bypasses stomatal immunity and addresses the contribution of water immunity. For syringe infiltration, we used mutants representing the extreme phenotypes from spray infection, *ht1-2, max2-4*, and *ghr1-3* (Figure 1D). We anticipated more bacterial growth in *ht1-2* than in the other mutants, as more closed stomata would allow higher apoplast hydration and water-soaking. However, after 24 hours post infection (HPI), we observed approximately two times less bacteria in *ht1-2* and *ghr1-3* compared to Col-0, while *max2-4* exhibited increased bacterial growth.

### Stomatal immunity is not the main determinant for the disease outcome

Successful *Pst* infection requires the bacteria to enter the plant through stomata and induce an aqueous phase inside mesophyll apoplast (Xin et al., 2016; Aung et al., 2018). Therefore, we first assessed the stomatal responses after *Pst* treatment (Figure 2, A and B). Stomatal aperture (width/height) of spray-inoculated plants was measured 1 hour after infection. All mutants had increased apertures in mock treated plants compared to Col-0, except *ht1-2*, which showed decreased aperture. *Pst* induced distinctive closure of stomata in Col-0 and, interestingly, in *ost1-3*, whereas the other mutants showed only mild or no response. Stomata of *ost1-3* was previously shown to be unresponsive to flg22 treatment (Zheng et al., 2018), but have a wildtype-like closure in response to SA and live *Pst* (Prodhan et al., 2018; Ou et al., 2022), suggesting that the initial pathogen-triggered stomatal closure is not dependent on OST1. These results suggests that impaired stomatal immunity cannot explain the disease outcome except for *max2-4*.

**Figure 2.**
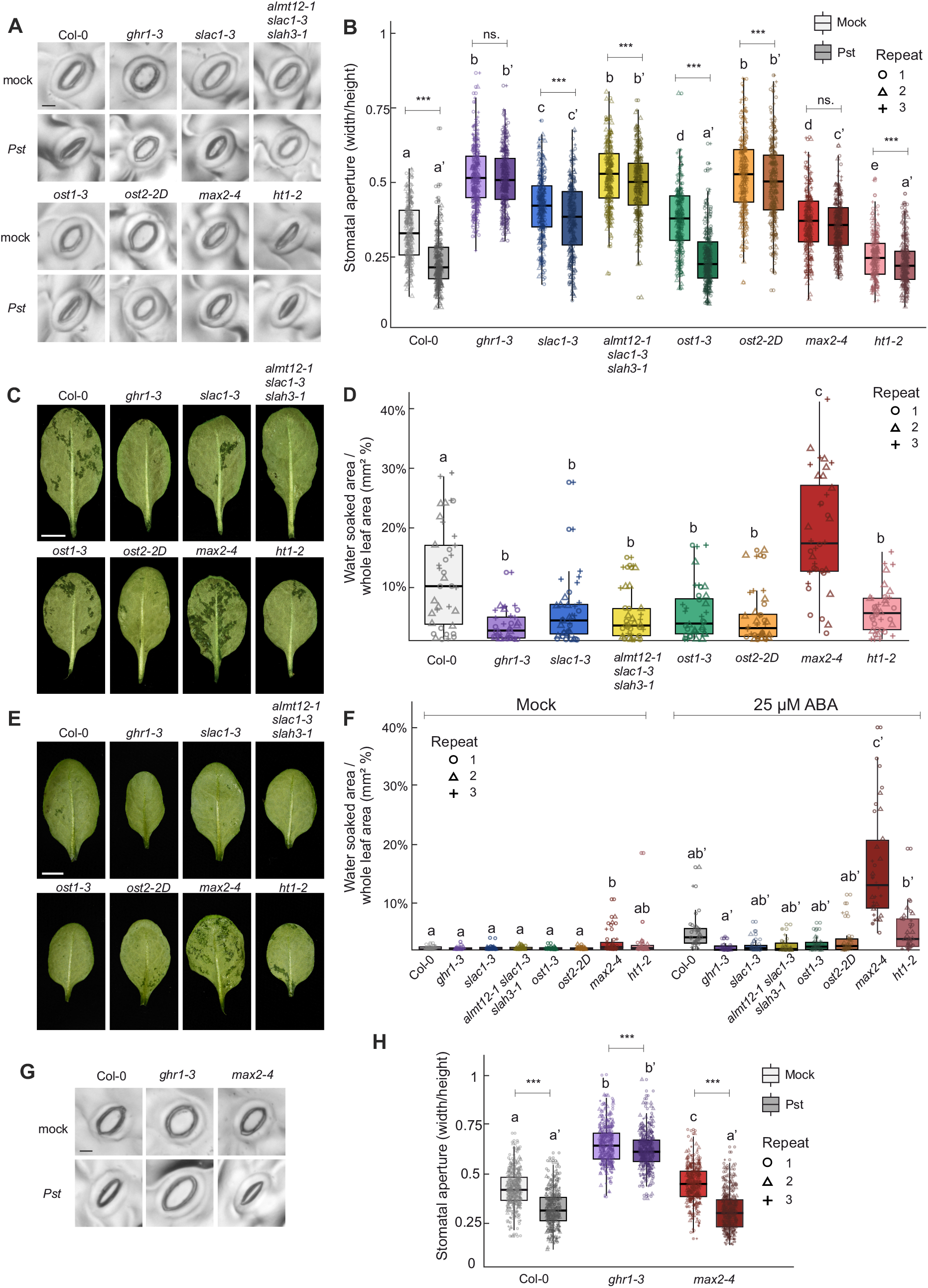
*Pst*-induced stomatal closure and *Pst*-and ABA-induced water-soaking in stomatal mutants. **A)** and **B)** *Pst*-induced stomatal closure in experimental lines. Plants were sprayed with mock or *Pst* (OD_600_= 0.2, 1 × 10^8^ CFU/mL), and epidermal imprints were taken 1 hour post-treatment. Representative images of stomata are shown (A), scale bar = 10 μm. Stomata from three plants per genotype across three independent experiments were analyzed (n ≈ 300 stomata in total per genotype). **C)** and **D)** *Pst*-induced water-soaking in stomatal mutants. Plants were sprayed with *Pst*, and abaxial leaf sides were photographed 24 HPI. The water-soaked ratio (%) was determined by dividing the water-soaked area by the total leaf area. Representative images are shown (C), scale bar = 5 mm. Two leaves from five to six plants per genotype across three independent experiments were analyzed (n = 36 leaves in total per genotype). **E)** and **F)** ABA-induced water-soaking in stomatal mutants. Plants were treated with 25 μM ABA or mock solution, and abaxial leaf sides were photographed 24 hours after treatment. Water-soaked area (%) was determined as above. Representative images of ABA-treated plants are shown (E), scale bar = 5 mm. **G)** and **H)** *Pst*-induced stomatal closure 24 HPI in Col-0, *ghr1-3*, and *max2-4*. Plants were sprayed with *Pst* as before, and stomatal imprints were taken 24 hours post-infection. Stomatal measurements (n ≈ 300 stomata in total per genotype/treatment) were obtained from two leaves per plant, four plants per genotype across three independent experiments. In (B), (D), (F), and (H), box plots represent three independent repeats with medians, interquartile ranges, and whiskers showing 1.5× interquartile range. A linear mixed-effects model followed by Tukey’s posthoc test was applied to evaluate statistical differences between genotypes. In (B) and (H), the statistical tests were conducted separately for mock and *Pst*-treatment (marked with apostrophe) groups. The effect of treatment was also evaluated, with statistical significance (p < 0.05) indicated by ***, and ns. representing non-significance.

### Water-soaking capacity dictates the disease outcome

To assess water immunity, we investigated the capacity of *Pst* to induce water-soaking (Figure 2, C and D). We sprayed plants with *Pst* and measured the water-soaked areas in leaves 24 HPI and calculated the afflicted area as a percentage of the whole leaf area (Figure 2D). The degree of water-soaking was elevated in *max2-4*, whereas it remained lower than Col-0 in all other mutants, reflecting the observed bacterial counts (Figure 1B). The disease outcome could thus be explained by water-soaking capacity.

We next evaluated the water-soaking capacity of the mutants independent of *Pst* to determine whether the observed water-soaking was a consequence or a contributing factor to disease outcome. ABA combined with high humidity was used as an alternative approach to induce water-soaking. ABA and high humidity, but also high humidity alone, induced elevated water-soaking in *max2-4* (Figure 2, E and F). A trend of ABA-induced water-soaking was also apparent in Col-0 and *ht1-2*, but not in other mutants. This suggested that the water-soaking capacity, likely associated with stomatal status, was a contributing factor to disease outcome.

*Pst* infection first leads to stomatal closure followed by re-opening at later stages of invasion to establish water immunity. To test if the water-soaking capacity (Figure 2, C and D) was associated with stomatal status at later stages of infection, we sprayed mutants representing the extreme phenotypes of water-soaking, *max2-4*, and *ghr1-3*, with *Pst* and measured stomatal aperture 24 HPI (Figure 2, G and H). The stomata of *max2-4* showed prominent closure at 24 HPI compared to mock treatment, whereas *ghr1-3* stomata remained open. Thus, the susceptibility of *max2-4* against *Pst* could be due to the combination of higher bacterial entry at the first step of infection (lack of stomatal immunity) (Figure 2B), followed by stomatal closure at later stages (lack of water immunity) (Figure 2H). For *ghr1-3*, the resistance is likely due to enhanced water-immunity that is overriding the lack of stomatal immunity.

### Expression of immunity and abiotic stress marker genes are not altered in stomatal mutants

To test if mutants had pre-activated defenses before bacterial infection, we used real time quantitative PCR (qPCR) to test marker genes for SA (*ICS1-ISOCHORISMATE SYNTHASE1*, a key gene in SA biosynthesis), JA (*JAZ1 - JASMONATE-ZIM-DOMAIN PROTEIN1*, involved in JA signaling, and *LOX4 - LIPOXYGENASE4*, a component of JA biosynthesis), ABA (*NDEC3 – NINE-CIS-EPOXYCAROTENOID DIOXYGENASE3*, an ABA biosynthesis gene), and abiotic stress (*P5CS1 - DELTA1-PYRROLINE-5-CARBOXYLATE SYNTHASE1*, involved in catalyzing the rate limiting step in proline biosynthesis) (Figure 3). Plants were grown as for *Pst* growth and stomatal experiments, and fully expanded leaves were used for qPCR. No significant changes in gene expression were found apart from the JA marker *LOX4*, which had higher expression in *max2-4*. We conclude that the tolerance to *Pst* infection observed in *ht1*-*2, ghr1-3, ost2-2d, slac1-3, ost1-3* and *almt12-1 slac1-3 slah3-1* cannot be explained by these mutants having pre-activated defenses.

**Figure 3.**
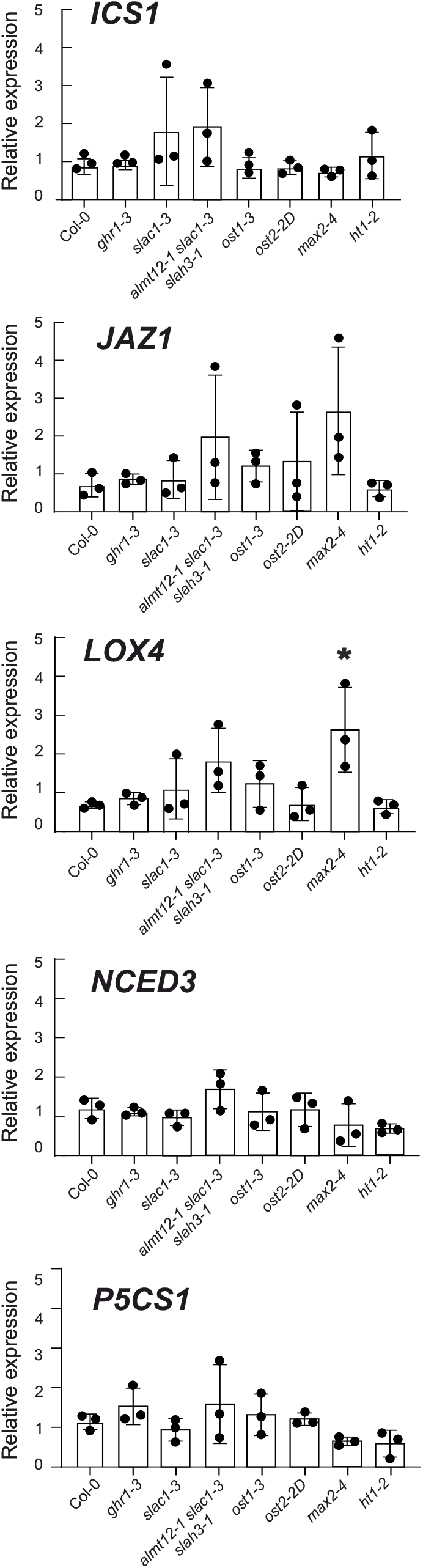
Expression of defense marker genes. Real-time quantitative PCR (qPCR) was used to assess the relative expression of *ICS1, JAZ1, LOX4, NCED3, P5CS1* in stomatal mutants and normalized against reference genes. Plants were grown under same conditions as for other experiments. Data represents three biological replicates, analyzed using one-way ANOVA followed by Tukey’s multiple comparisons test. Error bars indicate standard deviation, with significance (p < 0.05) marked by an asterisk.

## Discussion

Susceptibility of stomatal mutants was long attributed to the disrupted stomatal immunity and enhanced pathogen entry (Supplemental Table S1). However, recent evidence indicates that enhancing transpiration and stomatal openness can improve plant resistance in both Arabidopsis and rice (Zhang et al., 2019; Liu et al., 2022; Lajeunesse et al., 2023). With the recent advances in understanding the dynamics of plant disease and stomatal regulation, we aimed to clarify the relative importance of stomatal versus water immunity responses for plant resistance. Our results suggest that despite disrupted initial stomatal immunity responses, most mutants with more open stomata displayed significantly greater resistance against *Pst* compared to Col-0, supporting recent findings (Roussin-Léveillée et al., 2022; Hu et al., 2022; Liu et al., 2022; Lajeunesse et al., 2023). This resistance was observed regardless of whether the dysfunction affects stomatal closing or opening; as mutants with disrupted closure, e.g., inactive SLAC1, as well as the mutant with constitutive opening due to hyperactive H+-ATPase OST2, were all more resistant to *Pst* (Figures 1 and 2). Our results suggest that stomatal opening at post-invasive stages is crucial for plant resistance in a more general manner than previously recognized and can counteract the lack of stomatal immunity (Figure 4).

**Figure 4.**
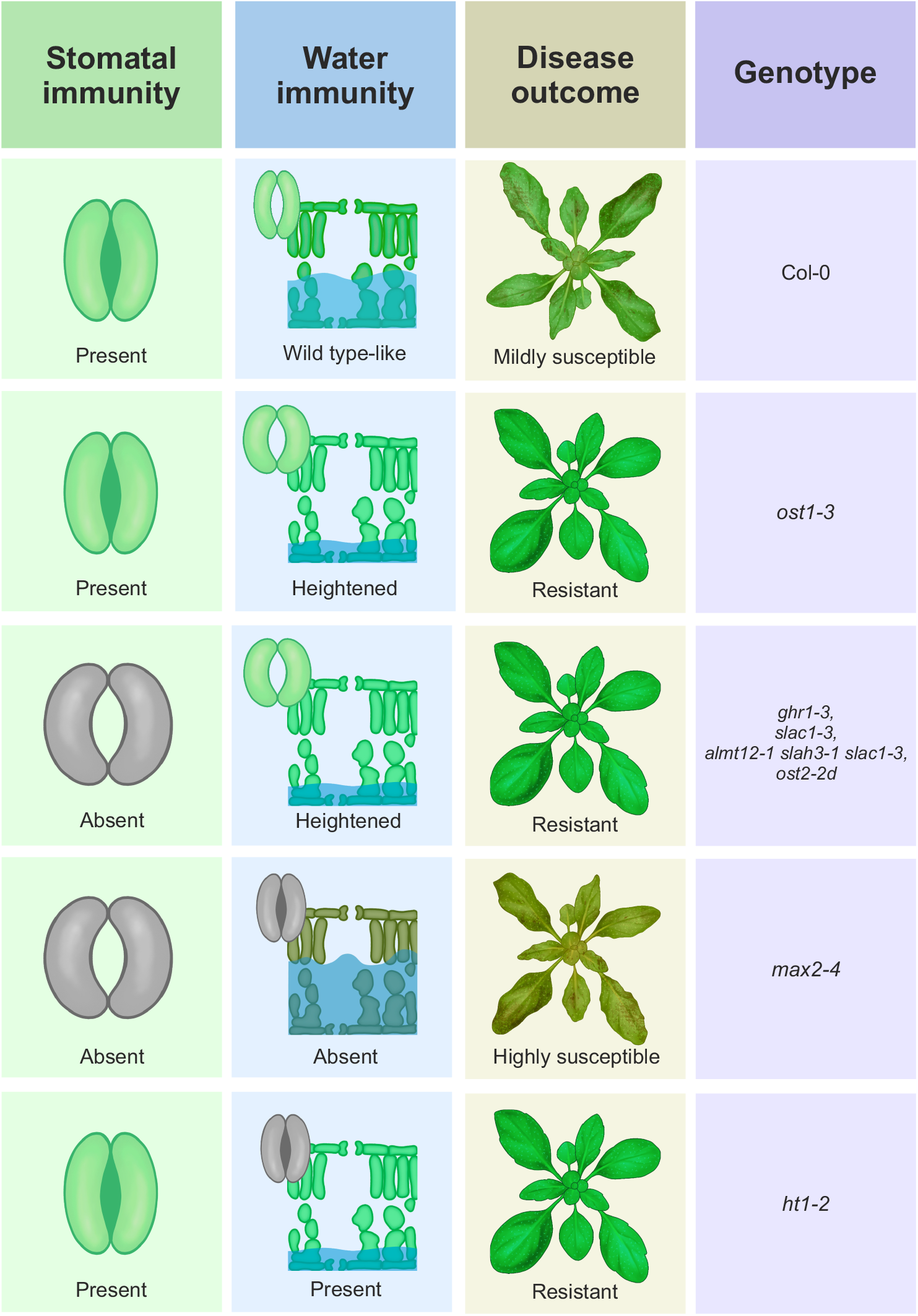
Model illustrating the relationship between temporal stomatal behavior, apoplastic hydration and disease outcome. Stomatal immunity (the ability to close stomata in response to pathogens) and water immunity (stomatal reopening and apoplastic hydration) are shown alongside disease outcomes for different genotypes. Dysfunction in stomatal behavior is indicated with grey color, showing the absence of functional closure or reopening. Water level (in blue) indicates water-soaking capacity. Water immunity is the main determinant of disease outcome and can override the absence of stomatal immunity.

As a mutant with constitutively closed stomata, we hypothesized that *ht1-2* would be more susceptible due to easier formation of an aqueous phase. However, *ht1-2* was more resistant against *Pst* after both spray- and syringe-infiltration (Figure1, B and D). The mechanisms of *ht1-2* resistance are unclear, but our study indicates they are not directly related to its stomatal phenotype. Interestingly, *ht1-2* exhibited similar degree of water-soaking to Col-0 after ABA treatment despite the higher resistance against *Pst* (Figure 2F). This suggests that the resistance against *Pst* in *ht1-2* could be due to other unexplored mechanisms, for example, mesophyll properties or heightened non-transcriptional immunity responses, as *ht1-2* didn’t show pre-activated defenses at the gene expression level (Figure 3).

The *max2-4* mutant exhibited a distinct susceptibility to *Pst* compared to other lines (Figure 1, B and C). Heightened susceptibility of *max2-4* was previously attributed to the lack of stomatal immunity (Piisilä et al., 2015). However, as the other open-stomata mutants exhibited heightened resistance, disrupted stomatal immunity cannot explain the susceptibility of *max2-4* alone. We propose that *max2-4* has impaired immunity at both steps of stomatal closure and reopening, allowing more *Pst* entry early on and heightened capacity to form higher water-soaking at later stages (Figure 4). Notably, *max2-4* exhibits a wild-type-like decrease in stomatal conductance following ABA treatment (Piisilä et al., 2015). We also observed stomatal closure 24 hours after *Pst* infection (Figure 2H), possibly due to the promoted ABA signaling by *Pst* (De Torres-Zabala et al., 2007; Hu et al., 2022; Roussin-Léveillée et al., 2022). Moreover, ABA alone induced a higher degree of water-soaking in *max2-4* compared to other lines (Figure 2F). Increased water accumulation of *max2-4* could thus be due to sensitivity to ABA signaling and/or other, e.g., anatomical properties.

The mutants *ht1-2* and *max2-4* further highlight the importance of experimental design in pathogen growth assays. For example, if only spray infection was used or a single time point early in the infection, it could be possible to speculate from our results that the resistance of *ht1-2* is due to its closed-stomata phenotype, while the susceptibility of *max2-4* is related to its disrupted stomatal immunity responses. Our study underscores how the temporally varying nature of *Pst* pathogenesis can easily lead to misunderstandings, often due to the oversights in experimental design. Previously, the importance of stomatal immunity has been emphasized in literature, potentially leading to inaccurate conclusions about mutant susceptibility.

Accurate reporting of experimental design and methods is also crucial. For example, it is well known that maintaining high humidity is important for successful *Pst* pathogenesis, often achieved by adding a sealed cover on infected plants (Katagiri et al., 2002). Indeed, elevated humidity levels directly correlate with increased *Pst* proliferation inside plant tissues (Xin et al., 2016). Furthermore, light also has a role in controlling plant immunity both on stomatal and whole-plant level; for example, it was shown that light can mitigate *Pst* infection possibly by enhancing SA signaling (Lajeunesse et al., 2023). However, increased light intensity can also reopen stomata, thus decreasing water-soaking (reviewed by Inoue & Kinoshita, 2017).

Here we have shown that stomatal dysfunction mutants exhibit increased resistance to *Pst* despite impaired initial closure of stomata. Our findings highlight the importance of water immunity in plant resistance against *Pst* in determining the disease progression (Figure 4). Although recent articles have suggested that water immunity (i.e., the prevention of aqueous space) is controlled by stomatal status, direct experimental evidence is scarce. Our study shows that stomatal regulation plays a significant role in managing apoplastic water status and, consequently, plant resistance.

## Materials and Methods

### Plant Material and Growth Conditions

Arabidopsis lines and their origins are listed in Supplemental Table S2. Plants were grown in 1:1 mixture of peat and vermiculite in chambers (Fitotron® SGC120, Weiss Technik) under 12 h light (170 μmol m−2 s−1)/12 h dark cycle, 23°C/19°C (day/night), 60%/70% relative humidity. For all experiments, 3.5-week-old plants were used.

### Bacterial Growth Assays

*Pseudomonas syringae* pv. *tomato* DC3000 was cultured overnight at 28 °C in liquid LB, diluted 1:50 in fresh LB, and cultured for 12 h. Bacteria was collected by centrifugation at 4000 rpm for 2 min and resuspended in 10 mM MgCl_2_, followed by two identical wash steps. Cell density was adjusted to OD600nm = 0.2 (1×10^8^ CFU/mL) or 0.02 (1×10^7^ CFU/mL) in 10 mM MgCl_2_ for spray and syringe inoculation, respectively, and 0.02% (v/v%) of Silwet L-77 was added.

For spray-infections, adaxial leaf surfaces were saturated with an airbrush spray. For syringe-infections three fully expanded leaves (leaf number 5-7) were infiltrated with a 2 mL needleless syringe until saturation. Infections were performed at mid-day. Trays were covered with plastic domes two hours prior to infections and again after the infections to maintain high humidity.

For bacterial counting, three leaves per plant were excised, surface-sterilized in 70% ethanol and washed twice in sterile milliQ. One leaf disc from each leaf was collected with 0,7 cm cork borer and ground in 10 mM MgCl_2_. Serial dilutions were plated on LB-Miller-plates and incubated at RT for ∼40 hours. Colonies were counted to determine colony-forming units (CFU). In each repeat, six plants were sampled, and the experiment was repeated three times.

### Water-soaking Assays

*Pst* spray-inoculation was conducted as above. For ABA spraying, 25 uM ABA with 0.02% (v/v%) Silwet L-77, and 0.1% ethanol in sterile milliQ were used. Same solution without ABA was used as mock treatment for ABA. Plants were sprayed at mid-day and trays covered with plastic domes to maintain high humidity. At 24 h post-spraying, two leaves per plant were excised and the abaxial sides were photographed. For each genotype and treatment, 5-6 plants were photographed per experiment and the experiment was repeated three times. Water-soaked area was measured using ImageJ and calculated as described by Wu et al., 2023. The water-soaking area was divided by total leaf area (mm^2^) to obtain water-soaked ratio (%).

### Stomatal Assays

*Pst* spray-inoculation was conducted as above. At indicated times, two leaves (leaf number 5-7) per plant were excised and a dental resin mixture (Xantopren M Mucosa; Kulzer) was applied on the abaxial side. After solidifying, the impressions were removed, and a cast of the epidermis was created with clear nail varnish. Nail varnish was removed with clear tape and applied on microscopy slide. Images were acquired with Leica DMLB microscope using Leica 20x Fluotar objective, and stomatal aperture ratio (width/height) was measured using ImageJ. Sampling of leaves was done in a randomized order, and acquisition of images and measurements were performed blind.

### RNA isolation and real time quantitative PCR

RNA was isolated using Spectrum Plant Total RNA Kit (Sigma-Aldrich). Four leaves (leaf number 5-7) were pooled together from four plants. Extracted RNA (2.5 μg) was treated with DNase I (ThermoFisher Scientific) and used for cDNA synthesis with Maxima Reverse Transcriptase (ThermoFisher Scientific). After cDNA synthesis, final volume was diluted to 100 μl, and 1 μl used for qPCR with 2X FastDye qPCR mix (KleverLab). Reactions were run on CFX Opus 384 System (Bio-Rad). Primers used for qRT–PCR, and their amplification efficiency are listed in Supplemental Table S2. Analysis of qPCR results was performed using the qBase3.4 program (CellCarta), (Hellemans et al., 2007). Three reference genes (PP2AA3, TIP41, YLS8) were used for normalization. Expression stability of reference genes (M-value) was evaluated with geNorm in qBase (Supplemental Table S2). Statistical analysis of qPCR data using three biological repeats was done with 1-way ANOVA and Tukey multiple comparisons test (GraphPad 10.2.3).

## Statistical analysis

Statistical analyses were conducted with R version 4.4.1 (R Core Team, 2022, https://www.R-project.org). Linear mixed-effect models with biological repeats as random grouping factor were fitted using function lmer from package lme4 (Bates et al. 2015). Pairwise-comparisons (Tukey’s method) were performed using emmeans package (Lenth et al. 2022).

## Acknowledgements

We thank Ville Pennanen for guidance with *Pst* assays.

## Author contributions

J.K., M.B., and M.S. conceived and designed the research. J.K., M.P., and M.B. performed research. J.K., S.E., M.B., and M.S. analyzed data. J.K., M.B. and M.S. wrote the manuscript. All authors discussed and interpreted the data and edited and commented the manuscript.

## Supplemental data

Supplementary Table S1. Literature showing discrepancies in mutant resistance against *Pst* and growth conditions.

Supplementary Table S2. Mutant lines and genotyping primers used in this study.

Supplemental Table S3. qPCR primers and expression stability of reference genes.

